# m^6^A methylation regulates RNA axonal localisation and translation in developing neurons

**DOI:** 10.1101/2025.09.01.671701

**Authors:** Braulio Martinez De La Cruz, Simon Galkin, Shiqi Wang, Yuxuan Jing, Subramanian Raju Nurani, Pierre-Luc Germaine, Gerhard Schratt, Antonella Riccio

## Abstract

Methylation on adenosine N6 (m^6^A) is an abundant post-transcriptional modification of the RNA that regulates almost the entire lifespan of RNA transcripts, from splicing and nuclear export to RNA stability and translation. Peripheral localisation of RNA is an event common to most cells and especially relevant in neurons where transcripts are trafficked to subcellular compartments to promote growth and differentiation in axons, and synaptic functions in dendrites. Here we show that in developing sympathetic neurons, m^6^A modification regulates RNA transport and local translation in axons. Critically, Nerve Growth Factor (NGF), a neurotrophin essential for axon growth and neuronal survival, promotes the axonal localisation of methylated transcripts and inhibits protein synthesis of trafficking RNAs, preventing premature protein expression. Noticeably, the translation of the bifunctional mRNA *Trp53Inp2* depended on the methylation of the 3*’*UTR, further supporting the key role of m^6^A modification in determining mRNA fate. Mutation of *Trp53Inp2* m^6^A elements resulted in ectopic translation, defects of axon extension and neuronal death. Together, these data show that m^6^A is critical for RNA peripheral localisation and spatial regulation of protein expression.

## Introduction

Neurons are structurally complex cells that maintain their extraordinary architecture at least in part, by compartmentalising gene expression^1^. Asymmetric localization of RNA is an evolutionarily conserved mechanism that allows spatial restriction of protein synthesis to cellular compartments^2,3^. In central and peripheral neurons, transcripts are transported to dendrites and axons^4-8^ where they are rapidly translated in response to extracellular cues, such as the neurotrophin Nerve Growth Factor (NGF)^9-13^. Despite initial difficulties in isolating RNA and ribosomes in axons, several studies have now demonstrated that hundreds of transcripts and atypical ribosomal units can be detected^14-16^.

In addition to the coding sequence that represents the blueprint for protein synthesis, most transcripts carry additional information in their 5*’* and 3*’* untranslated regions (UTRs) that is essential for RNA localisation and translation^13,17-20^. RNA localization depends on the interaction of RNA-binding proteins (RBPs) with the cis-acting elements mostly found in the 3*’*UTRs of the localised transcripts^21^. The binding of RBPs to RNA, however, does not depend uniquely on the sequence of the RNA and relies on the secondary and tertiary structure, making the identification of localisation elements difficult^22^.

Akin to DNA, RNA can be epigenetically modified at many residues. Over one hundred modifications of the RNA have been identified so far, and many are linked to specific aspects of RNA metabolism, from RNA splicing and nuclear export to transport and translation^23,24^. N^6^-methyladenosine (m^6^A), one of the most common epigenetic modifications of RNA, was discovered over 50 years ago. It is especially abundant in neurons where it is enriched in conserved regions of the 3*’*UTRs and near stop codons of transcripts regulating neuronal development and differentiation^25-27^. Like the epigenetic modifications of chromatin, m^6^A may change the affinity and specificity of RBPs binding, thereby regulating nearly every aspect of the metabolism of RNA. m^6^A elements are recognised by specific RBPs named readers that mediate RNA splicing, stability and translation^24,28^. Recent studies have shown that RBPs involved in neurodegenerative disorders such as TAR DNA-binding protein 43 (TDP43) and Fragile X Mental Retardation Protein (FMRP) bind to methylated RNA in Amyotrophic Lateral Sclerosis (ALS)^29-31^ and Fragile X syndrome patients respectively^32,33^, suggesting that m^6^A may play a pivotal role in these disorders. Despite these observations however, the role of m^6^A in regulating RNA metabolism during neuronal development and in controlling RNA transport and translation in dendrites and axons remains largely unknown.

Here we show that in developing sympathetic neurons, exposure to the survival and growth-promoting neurotrophin Nerve Growth Factor (NGF)^34,35^ induces the rapid transport of m^6^A-modified transcripts in axons and is essential for axon extension. m^6^A sequencing performed on RNAs isolated from either cell bodies or axons revealed that methylated RNAs rapidly accumulate in axons in response to NGF, remaining largely unchanged in cell bodies. Importantly, Translating Ribosome Affinity Purification (TRAP)^36^ followed by sequencing revealed that in response to rapid stimulation with NGF, methylated RNAs are mostly untranslated in axons. Analysis of *Trp53Inp2*, a bifunctional mRNA that is highly methylated and not translated in sympathetic neurons^37^, revealed that m^6^A modification of three adenosines within the 3*’*UTRs is essential for *Trp53Inp2* transport and translation. Moreover, an *in vitro* transcribed non-methylatable *Trp53Inp2* transcript was unable to rescue the axon growth defects observed in *Trp53Inp2* mutant mice, indicating that m^6^A modification is essential for *Trp53Inp2* functions. Together, these data uncover a novel mechanism regulating RNA trafficking in neurons and reveal m^6^A modification as an essential mechanism for mediating NGF-dependent axon growth.

## Results

### NGF mediates the localization of m^6^A-methylated RNA in sympathetic neuron axons

Sympathetic neurons require NGF for survival, growth and axon extension during development. We previously discovered that NGF-dependent localisation of RNAs in axons is essential for maintaining axon extension and to promote cell survival^12,13^. First, we asked whether NGF regulates transcript methylation in sympathetic neurons. Neurons were cultured with NGF and after 5 days NGF was withdrawn for 18 hours followed by restimulation with NGF for 1 hour. Dot blot analysis of m^6^A methylated RNA indicated that removal of NGF resulted in increased RNA methylation when compared to both control and restimulated neurons (**Fig.1A**). In most cells m^6^A is added co-transcriptionally by a methylation complex including the methyltransferase METTL3, METTL14 that stabilizes METTL3/METTL14 interaction, and the Wilm*’*s Tumour 1 Associated Protein (WTAP) that is necessary for METTL3 import to the nucleus^38,39^. The intracellular signals regulating the assembly and the activity of the RNA methylase complex are mostly unknown however, all proteins are phosphorylated at several residues^40^ and METTL3 phosphorylation at S^43^ is known to stabilise and increase METTL3 methyl-transferase activity^41^. Western blot analysis indicated that NGF withdrawal increased METTL3 phosphorylation (**Fig.1B, C**), suggesting that phosphorylation of METTL3 may play a role in mediating NGF-dependent m^6^A methylation. Given that under these conditions, axon growth is severely stunted, a possible interpretation of these results is that NGF withdrawal for a short time increases the rate of m^6^A methylation of RNA transcripts that remain mostly stored in cell bodies.

**Fig. 1.**
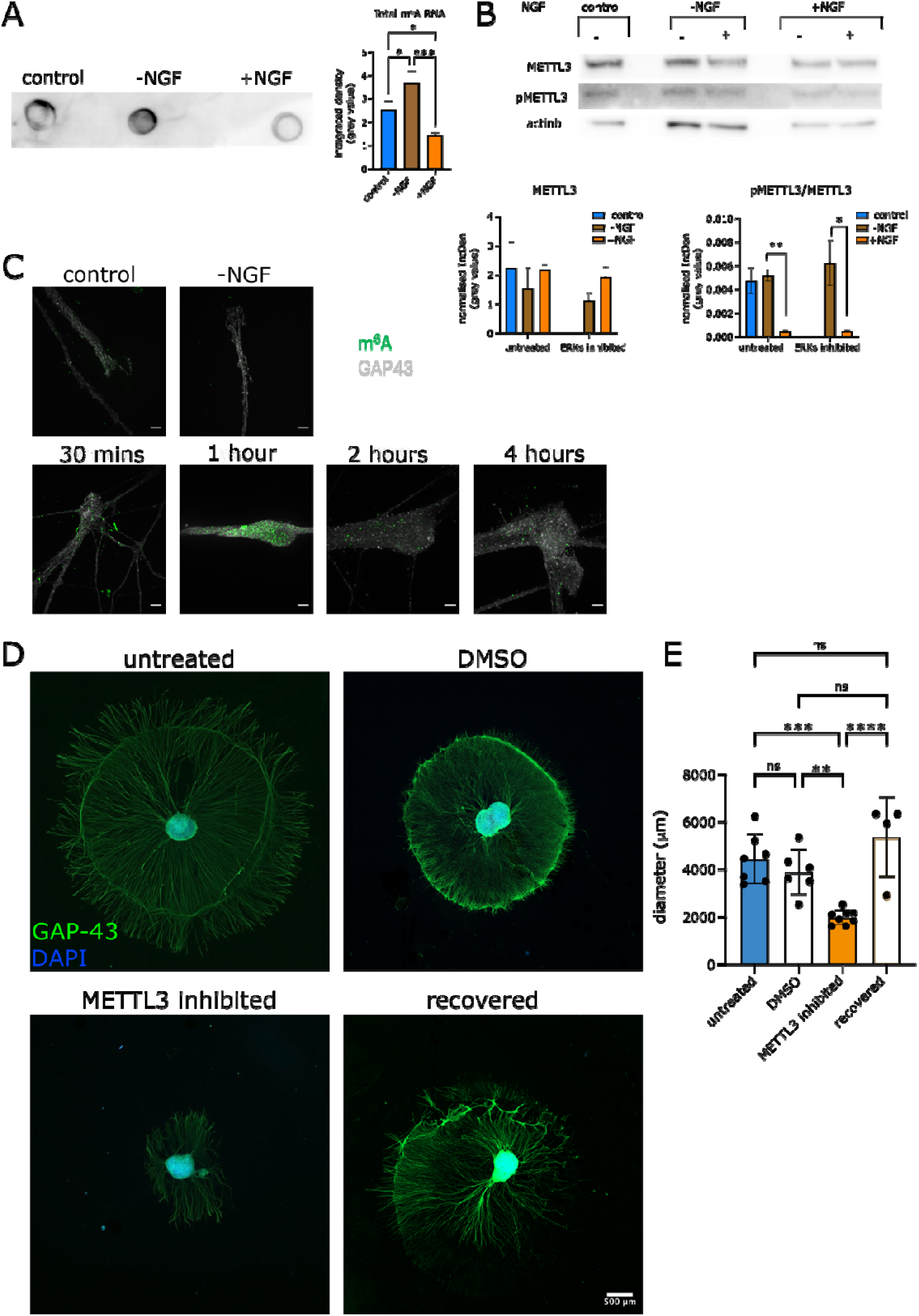
m^6^A-methylated RNA is transported to sympathetic neuron axons. (**A**, *left*) RNA dot blot of m^6^A-methylated mRNA under the indicated NGF condition. (*Right*) Quantification of dot pixel value intensity Ordinary one-way ANOVA and Tukey’s multiple comparison test: Control F vs -NGF *p*=0.0256; Control vs +NGF *p*=0.0288; -NGF vs +NGF *p*=0.0009. n = 3 (**B**) Western Blot of sympathetic neurons in control, -NGF, or +NGF conditions (see methods). All lanes contained approximately 30 μg of proteins. *Below*: Quantification of total METTL3 and pMETTL3 bands. pMETTL unpaired t-test: Untreated-NGF vs untreated+NGF *p*=0.0019, inhibited -NGF vs inhibited+NGF *p*=0.0346. All other comparisons non-significant. n=3 (**C**) m^6^A Immunofluorescence of methylated RNA in distal axon tips of sympathetic neuron explants under the indicated conditions (see methods). Scale bar=10 μm. (**D**) Sympathetic neuron explants grown for 5 days in NGF (control) and exposed to either the METTL3 Inhibitor STM2457 or vehicle (DMSO). In parallel experiments, the METTL3 inhibitor was removed after 5 days, and explants were cultured for 3 more days (recovered). Scale bar=500 μm. (**E**). Quantification of D. Ordinary one-way ANOVA test was performed. NS = non-significant. Untreated vs Inhibited: *p*=0.0004. DMSO vs Inhibited: *p*=0.0075. Inhibited vs Recovered: *p*< 0.0001. n=4-7.

To study whether m^6^A methylation regulates transcripts localisation in axons we cultured early postnatal sympathetic cervical ganglia (SCG) with NGF. After 4 days, NGF was withdrawn and the caspase inhibitor BAF was added to prevent cell death. SCG explants were then exposed to NGF for various times and subject to immunofluorescence to detect m^6^A methylated RNA transcripts. We found that both in control conditions, when neurons were maintained with NGF, and following NGF withdrawal the levels of m^6^A in axons were low (**Fig.1C**). Conversely, in response to NGF stimulation for 30 minutes, methylated transcripts rapidly accumulated to distal axons and growth cones. NGF stimulation for longer times revealed that m^6^A RNA methylation returned to basal levels within two hours, indicating that rapid transport of methylated transcripts represents an early response to NGF stimulation (**Fig.1C**).

In rodents, during the early stages of development, NGF is released in limited amounts by the target tissues to promote neuronal survival, axon extension and for maintaining axon integrity ^35^. To study whether RNA methylation was necessary for axon growth, SCG explants were exposed to the METTL3 Inhibitor STM2457^42^ for 4 days in the presence of NGF (**Fig. S1A**). We observed a striking reduction of axon growth when compared to untreated or vehicle-treated controls (**Fig.1D** and **E**). When STM2457 was removed, axon growth fully recovered to reach normal length, indicating that under these conditions, cell viability was largely conserved (**Fig. S1B**). Together, these results indicate that NGF induces the rapid transport of m^6^A-modified RNAs in axons and that this event is essential for NGF-dependent axon growth.

### The compartmentalised m^6^A epitranscriptome in developing sympathetic neurons

The finding that NGF induces a substantial accumulation of m^6^A modified RNAs in distal axons and growth cones prompted us to perform a genome wide analysis to identify methylated transcripts in both cell bodies and axons of sympathetic neurons. Neurons were grown on semi-permeable membranes (**Fig.2A**) with NGF (50 ng/ml) in the axonal compartment. After 5 Days, NGF was withdrawn, and neurons were maintained with the caspase inhibitor boc-aspartyl(OMe)-fluoromethylketone (BAF)^43^ for 18 hours to prevent cell death. Axons were then exposed to NGF for 1 hour, RNA was isolated from axons and cell bodies in all three conditions (control, -NGF or +NGF, n=4 per condition), and immunoprecipitated with m^6^A-specific antibody (**Fig.2A**). Libraries were prepared from both input and immunoprecipitated samples and subject to sequencing. Notably the m^6^A consensus motif RAAC was overrepresented in all groups. Unfortunately, NGF withdrawal substantially reduced RNA localisation in axons^13^, therefore the amount of RNA recovered from NGF-deprived axons was insufficient for ultra-low input sequencing. First, we analysed the distribution of m^6^A RNA transcripts in axons and cell bodies of neurons maintained with NGF in homeostatic conditions. Strikingly, ∼43% (3070 of the 6,819) transcripts were hypermethylated in axons, whereas only 595 transcripts (∼5% of 10,871) where found to be hypermethylated in cell bodies (**Fig.2B**). *Trp53Inp2*, a bifunctional mRNA that interacts with NGF receptor TrkA and mediates NGF signalling^37^, was also found to be among the methylated RNAs highly enriched in axons. Heatmaps revealed a remarkable change of methylated transcript localisation in axons following acute exposure to NGF, whereas fewer variations were observed in cell bodies (**Fig.2C**). RNAs encoding for *Tro, Trappc2, Grin3a*, and *Grin1* were enriched in axons in response to NGF, whereas transcripts encoding for *Slc2a, Fdxr* and *Snc1a* were depleted following stimulation. GO analysis confirmed that m^6^A methylated RNAs related to NGF signalling pathways and vesicle trafficking were enriched in axons whereas methylated transcripts regulating synaptic functions and cell metabolism were overrepresented in the cell bodies (**Fig.2D, E**). Finally, although studies on RNA methylation focus on mRNA, we also found that many non-coding RNAs were enriched depending on location and treatment. Examples included *Snord17* and *Ftx* (**Fig. S3**).

**Fig. 2.**
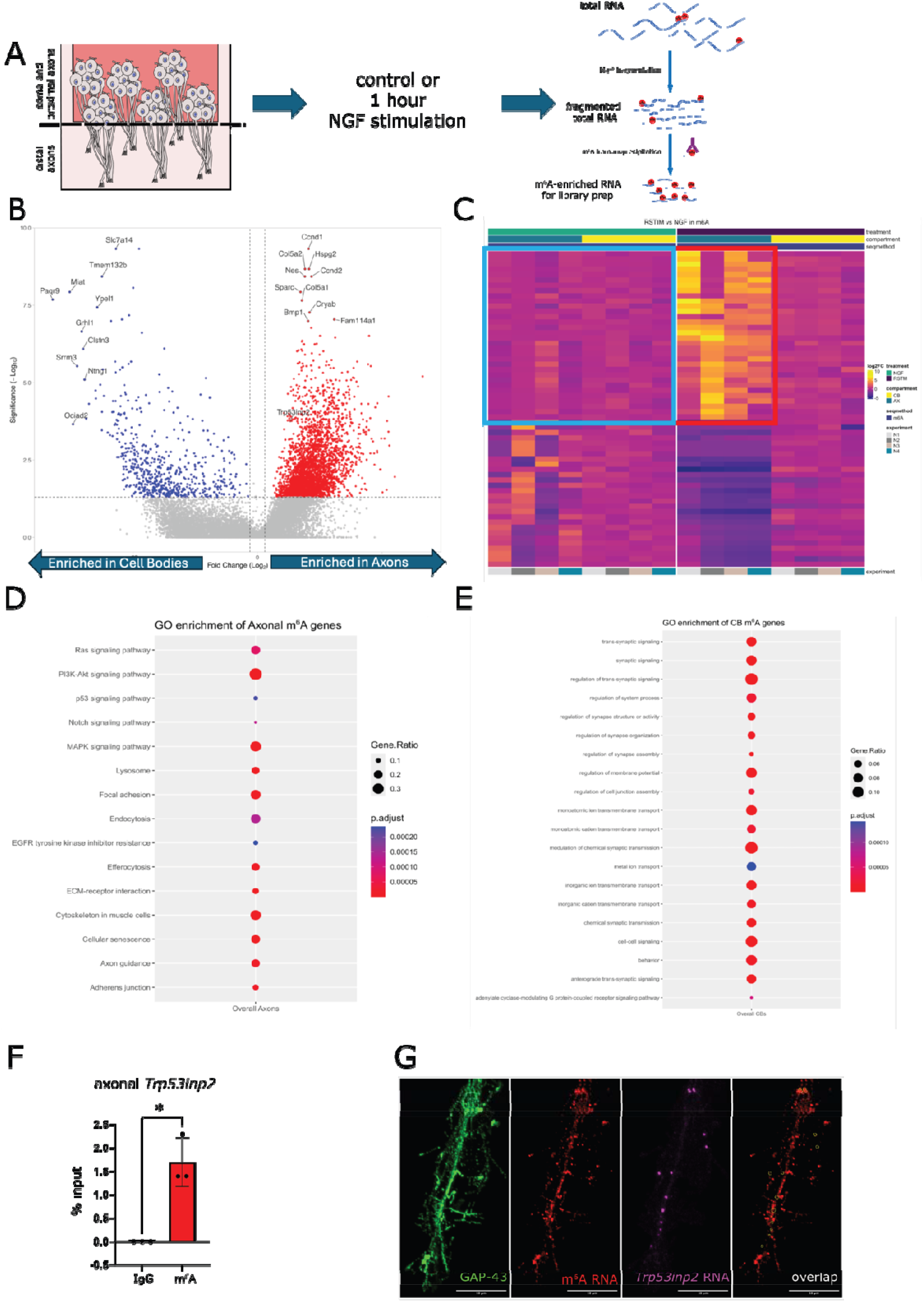
The m^6^A epitranscriptome in axons and cell bodies of sympathetic neurons. (**A**) Schematic representation of meRIP-seq in sympathetic neurons. Sympathetic neurons were cultured on semi-permeable membranes and total RNA from either the cell bodes or axons was subject to fragmentation and immunoprecipitation using an m^6^A-specific antibody. Libraries were prepared from both input and immunoprecipitated samples and sequenced. (**B**) Volcano plot of m^6^A-enriched RNAs in axons (3070) and cell bodies (595). (**C**) Heatmap showing the top 60 genes differentially methylated in control (*left*) and NGF stimulated conditions (*right*). (**D, E**) Gene ontology analysis of axonal genes enriched in axons (**D**) or cell bodies (**E**). (**F**) meRIP-RT-qPCR of axonal *Trp53inp2*. Unpaired t-test with Welch’s correction *p*=0.0034 (**G**) Immuno-FISH of m^6^A RNA (red) and *Trp53inp2* RNA (magenta) of sympathetic neuron axons. GAP-43 (green) was used to visualize the axons. Scale bar=10 μm. n=3

To validate some of the most enriched m^6^A methylated axonal transcripts, methylated RNA immune precipitation (meRIP)-RTqPCR and fluorescence *in situ* hybridization (FISH) combined with m^6^A immunostaining were performed. We confirmed that the *Trp53Inp2* transcript was significantly hypermethylated in axons (**Fig.2F**). Exposure to NGF for 1 hour increased co-localisation of m^6^A immunofluorescence and *Trp53inp2* RNA FISH in distal axons. Similar results were observed when other methylated RNAs highly enriched in axons, such as Map1b were analysed (**Fig.S4A**).

### Active transport of methylated transcripts in axons

Next, we asked whether the accumulation of m^6^A methylated RNA transcripts in axons was due to localised methyltransferase activity or to active transport from the cell bodies. Western blotting of proteins obtained from either cell bodies or axons indicated that METTL3 and METTL14 were not expressed in axons (**Fig.3A, B** and **Fig.S4B**), ruling out the possibility that transcripts may be methylated in axons in response to NGF and further confirming that m^6^A modifications are added co-transcriptionally in the nucleus^24^. Analysis of m^6^A immunostaining in axons exposed to NGF for various times confirmed that methylated RNA is rapidly transported to the most distal part of the axons, with levels of methylated transcripts returning to homeostatic levels within 2 hours (**Fig.3C, D**). Notably, in neurons maintained with NGF, m^6^A methylated RNAs was distributed homogenously along the axons, indicating that in homeostatic conditions methylated transcripts are constantly trafficked to axons.

**Fig. 3.**
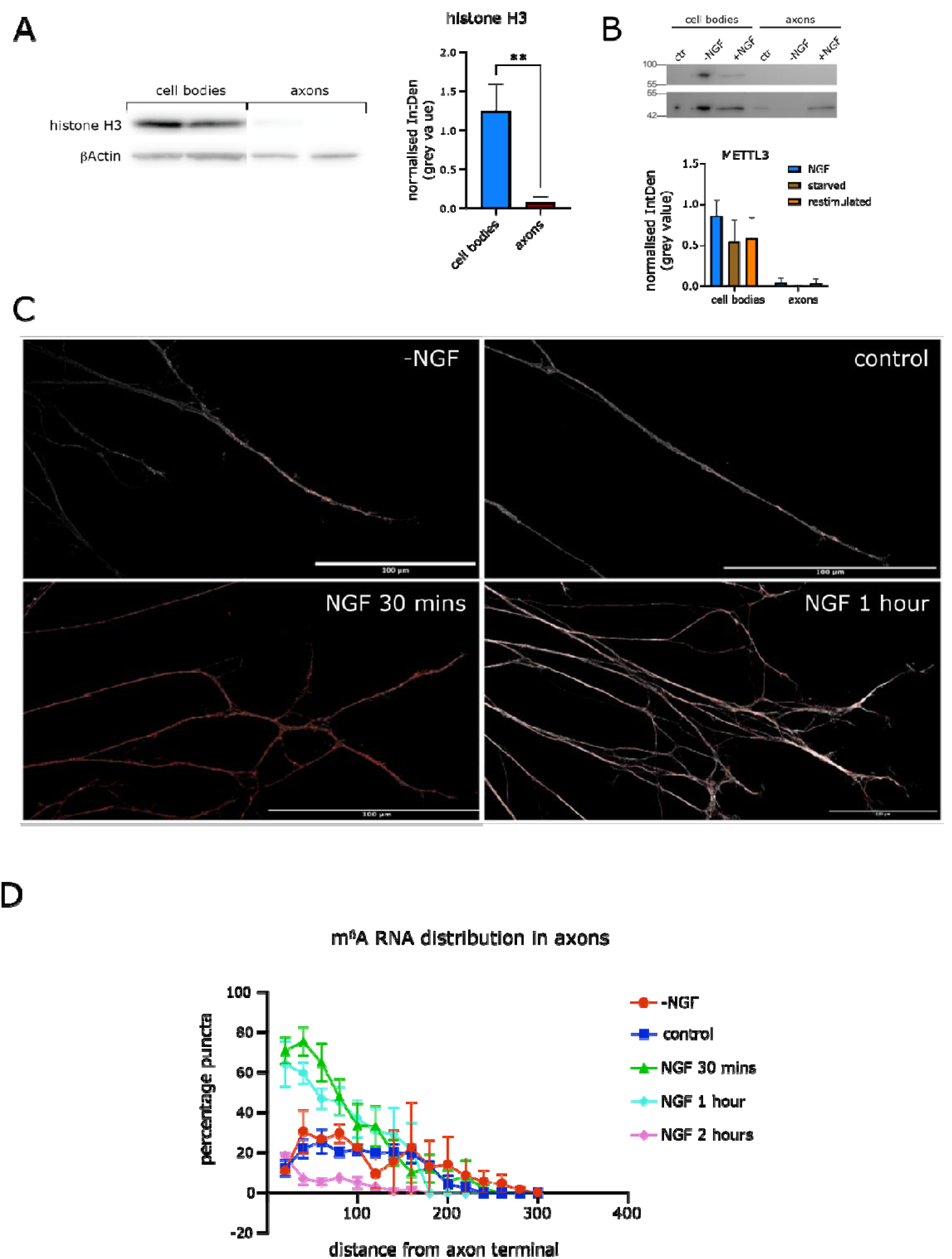
m^6^A-RNAs accumulate to distal axons in response to NGF. (**A**) Western Blot of cell bodies and axons (depicted in **Fig. 2A**) but not H3 histone protein. All lanes contained approximately 30 μg of total protein. (*Right*) Quantification of (**A**). Two-tailed unpaired t-test with Welch’s correction *p*=0.0055. n = 4 (**B**) METTL3 western blotting of cell bodies and axons from neurons in the indicated conditions. All lanes contain approximately 30 μg of total protein. (*Bottom*) Quantification of (**B**). (**C**) Immunostaining of m^6^A-RNAs (red) in distal axons of sympathetic neurons in the indicated conditions. GAP-43 in greyscale. Scale bar=100 μm. (**D**) Quantification of methylated RNA as a function of distance from the tip of the axons towards the cell bodies. Axon terminal =0. (n=10 for NGF 30 min, n=5 for all other conditions). Two-way ANOVA multiple comparisons test: -NGF vs 30 mins, *p*<0.0001; -NGF vs 1 hour, *p*=0.0412; +NGF vs 30 mins, *p*<0.0001, +NGF vs 1 hour, *p*=0.0393; 30 mins vs 2 hours, *p*=0.0001. All other comparisons non-significant.

### m^6^A methylation represses the translation of axonal mRNA transcripts

Compartmentalization of RNA is a ubiquitous process common to most cell types and detected across species^2^. In neurons, it serves multiple purposes. First, it allows the rapid expression of proteins in response to localised stimuli, generating local translation only where and when proteins are needed^7,13,14,44^. Second, it avoids delay and potential protein degradation associated with the cellular trafficking of RNAs and proteins along long distances^45^. Finally, it avoids ectopic expression of proteins in certain cellular compartments reducing potential toxicity. We had previously shown the *Trp53Inp2* is not translated in sympathetic neurons, and interacts with the NGF receptor TrkA, enhancing NGF-TrkA signalling^37^. *Trp53Inp2* belongs to a new class of bi-functional mRNAs that harbour both coding and non-coding functions^20,46^, and in other cell types is translated into a protein that regulates autophagy^47^. We took advantage of the bifunctional nature *Trp53Inp2* to study whether m^6^A methylation regulates translation in axons. Western blotting of the mouse myoblast C2C12, a cell lines that expresses Trp53Inp2 protein^37,47^, showed that differentiation into myotubes inhibited translation (**Fig.4A**), despite expressing similar mRNA levels (**Fig.4B**). Remarkably, *Trp53Inp2* methylation was much higher in differentiated C2C12 (dC2C12), compared to naïve cells (**Fig.4C**). To study whether inhibition of m^6^A methylation influenced *Trp53Inp2* translation, siRNA targeting METTL3 was transfected in dC2C12, and Trp53Inp2 protein was assessed by western blotting. siMETTL3 greatly decreased METTL3 expression when transfected in dC2C12 cells, compared to scrambled siRNA or a siRNA targeting GAPDH (**Fig.4D**). Next, full length, *in vitro* transcribed *Trp53Inp2* was transcribed either as an unmethylated transcript (0%) or with 33% or 100% adenosines replaced by methylated adenosines. Western blotting of cells transfected with *Trp53Inp2* methylated at various levels revealed that *Trp53Inp2* m^6^A methylation inversely correlated with translation (**Fig.4E**). When dC2C12 were transfected with siMETTL3, endogenous *Trp53Inp2* translation was remarkably increased irrespective of the *Trp53Inp2* transcripts transfected, indicating that m^6^A methylation acts as a translational switch between coding and non-coding functions.

**Fig. 4.**
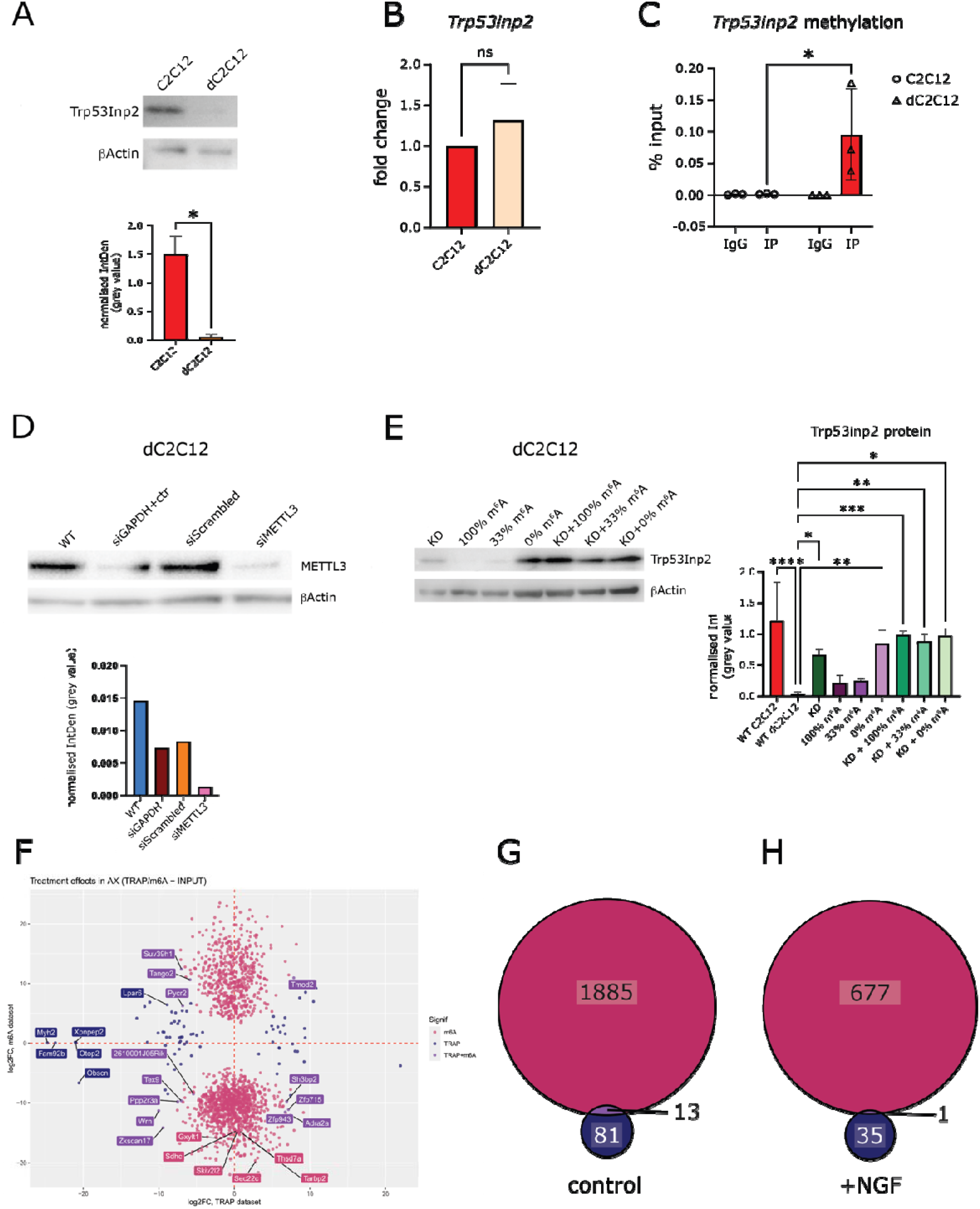
m^6^A methylation inhibits Trp53Inp2 translation. (**A**, *top*) Trp53inp2 western Blot of C2C12 and differentiated C2C12 (dC2C12) cells. All lanes contained approximately 20 μg of total protein. *(Bottom)* Quantification of A. Two-tailed unpaired t-test with Welch’s correction *p*=0.0026. n=4 (**B**) RT-qPCR of *Trp53inp2* expression in C2C12 and dC2C12 normalised to β-actin expression in C2C12 cells. Two-tailed unpaired t-test with Welch’s correction *p* = 0.3505. (**C**) meRIP-RT-qPCR of m^6^A-methylated *Trp53inp2*. Two-way ANOVA with Sidak’s multiple comparisons correction C2C12 IP vs dC2C12 IP *p*=0.0263. n=3 (**D**) Western Blotting of dC2C12 cells transfected with the indicated vectors. All lanes contained approximately 30 μg of total protein. (**E**) Western blotting of C2C12 and dC2C12 cells transfected with the indicated siRNAs and *in-vitro* transcribed *Trp53inp2* with 100%, 33% or 0% methylated adenosines. All lanes contained approximately 30 μg of proteins *Right*: Quantification of E. Ordinary one-way ANOVA *p*<0.0001. Dunnet’s multiple comparisons test of each condition vs WT dC2C12. WT C2C12 *p*<0.0001; KD *p*=0.0224, 0% m6A *p*=0.0029; KD+100% m6A *p*=0.0006; KD+33% m6A *p*=0.0019; KD+0% m6A *p*=0.0007. n=3 (**F**) Dot plot comparing fold-change of significantly enriched genes in axonal meRIP-seq (red) and axonal TRAP-seq (blue) datasets, as well as genes significantly enriched in axons in both datasets (purple). Top genes are annotated. (**G**) Venn diagram of the number of unique and overlapping enriched genes in axons of meRIP-seq and TRAP-seq. (**H**) Venn diagram of the number of unique and overlapping enriched genes in NGF restimulated axons in meRIP-seq and TRAP-seq.

To study the role of m^6^A methylation in regulating translation of the wider axonal transcriptome we performed Translating Ribosome Affinity Purification followed by RNA sequencing (TRAP-Seq)^36^. mRNAs associated with the ribosomal subunit Rpl22 expressed in either cell bodies or axons of sympathetic neurons were isolated and sequenced. Transgenic mice expressing HA tagged Rpl22 were crossed with DHB-Cre mice expressing Cre recombinase under the control of the sympathetic neuron specific dopamine β hydroxylase (*dbh*) promoter^48^. Sympathetic neurons were grown in compartmentalised chambers in conditions similar to the ones used for m^6^A-RIP sequencing. NGF was withdrawn for 18 hours and neurons were either restimulated for 1 hour or left untreated. Results were validated by Puro-PLA (proximity ligation assay; **Fig.S5A, B**). TRAP-Seq data revealed that the biggest changes occurred in axons, although only 94 RNAs are associated with ribosomes in axons after 1 hour of exposure to NGF (**Fig.S5C-G**). Surprisingly, we also discovered some non-coding RNAs, including *MALAT1* which has recently been found to translate a small peptide in neurons in response to membrane depolarisation^49^. The overlap of m^6^A methylated transcripts enriched in axons or in response to a short NGF stimulation with ribosome-associated mRNAs under similar experimental conditions revealed that m^6^A methylation did not substantially affect ribosome interactions (**Fig.4F-H)**. It should be noted that TRAP-Seq measures the engagement of RNA with ribosomes. Thus, a potential interpretation of these data is that at least some untranslated axonal mRNAs may be bound to ribosomes and maintained in a *“*poised*”* state without engaging in productive translation.

### Binding of TDP-43 regulates axonal localisation of methylated Trp53Inp2

The finding that NGF induces rapid transport of m^6^A methylated *Trp53Inp2* prompted us to study whether inhibition of METTL3 affected axonal RNA transport. SCG explants were grown for 5 days, NGF was withdrawn for 18 hours followed by re-exposure to NGF for 1 hour and *Trp53Inp2* was visualised in both cell bodies and axons using FISH (**Fig.5A, B**). As expected, NGF induced the accumulation of m^6^A methylated *Trp53Inp2* in distal axons (**Fig.5C**). Importantly, no changes were observed in *Lin7c*, a transcript chosen as a control for being methylated equally in all conditions and cell compartments (**Fig.S6A, B**). While the absolute number of *Trp53inp2* puncta increased after NGF stimulation, the methylation coefficient remained approximately 0.85 in control and restimulated axons (**Fig.5D**). Exposure to the METTL3 inhibitor STM2457 inhibited *Trp53Inp2* axonal transport in both control and NGF-stimulated neurons. Conversely, METTL3 inhibition had no effect on *Trp53Inp2* localisation in cell bodies and proximal axons, indicating that methylation does not play a key role in nuclear export and stability of the transcript (**Fig. 5A** and **Fig.S6C, D**).

**Fig. 5.**
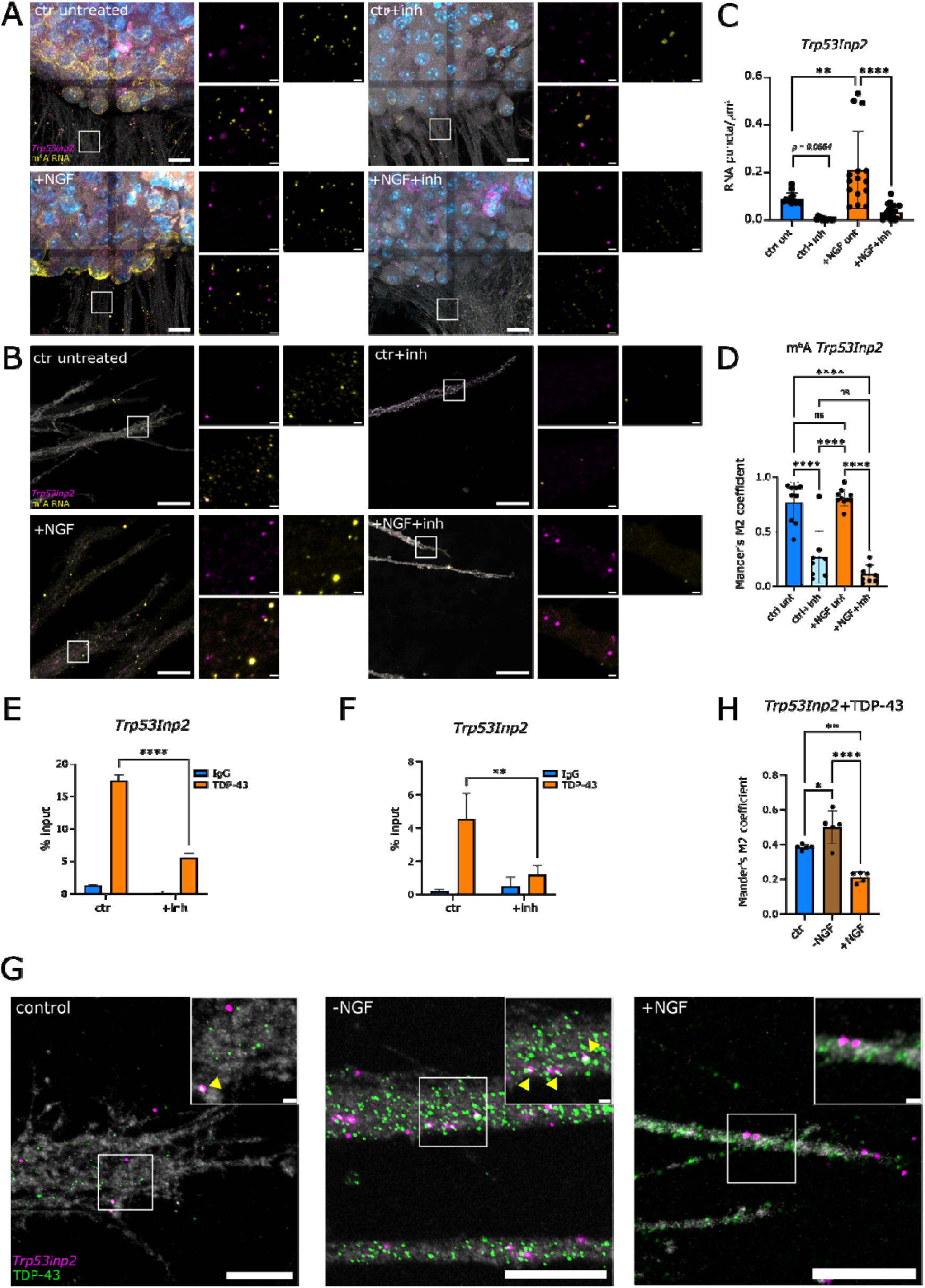
Axonal transport of methylated Trp53inp2 is mediated by TDP-43. Confocal microscopy images of *Trp53inp2* HCR RNA FISH (magenta) and m^6^A-RNA (yellow) in cell bodies and proximal axons (**A**) or distal axons (**B**) of sympathetic ganglia explants. Explants were treated with a combination of NGF and the METTL3 inhibitor, as indicated. Scale bar=10 μm. GAP-43 was used as a cytoskeletal marker (grey). Explants were treated with a combination of NGF and METTL3 inhibitor, as indicated. Scale bar=10 μm. (**C**) *Trp53Inp2* mRNA quantification of the data shown in (**A**), and normalised by axonal area. Ordinary one-way ANOVA p<0.0001. Tukey’s multiple comparisons tests. Ctrl unt vs ctrl inh p=0.0564; Ctrl unt vs +NGF unt p=0.0021; +NGF unt vs +NGF inh p<0.0001. n=13-19 (**D**) Quantification of methylated *Trp53inp2* shown in (**B**) as inferred by colocalization with anti-m^6^A immunofluorescence. Mander’s coefficient M2 quantified only sites with *Trp53inp2* signal. Ordinary one-way ANOVA p<0.0001. Tukey’s multiple comparisons tests: Ctrl Unt vs ctrl Inh p<0.0001; Ctrl unt vs +NGF unt p=0.9247; Ctrl unt vs +NGF inh p<0.0001; Ctrl inh vs +NGF unt p<0.0001; Ctrl inh vs +NGF inh p=0.2128; +NGF unt vs +NGF inh p<0.0001. n=8-10 (**E**) TDP-43/*Trp53inp2* RIP-RTqPCR of sympathetic neurons. Two-way ANOVA inhibition factor p<0.0001, antibody used p<0.0001, interaction p < 0.0001. Sidak’s multiple comparisons test ctr IP vs inh IP p< 0.0001. n = 3 (**F**) RIP-RTqPCR of TDP-43 interaction with *Trp53inp2* in distal axons of sympathetic neurons. Two-way ANOVA inhibition factor p=0.0144; Antibody used p< 0.0009; Interaction p=0.0062. Sidak’s multiple comparisons test ctr IP vs inh IP=0.0027. n=3 (**G**) Confocal microscopy of Immuno-FISH of *Trp53inp2* (magenta) and TDP-43 (green) in distal axons in the indicated conditions. Yellow arrows indicate examples of RNA-protein colocalization visualised in white. Scale bar=10 µm. Phalloidin 488 used as a cytoskeletal marker. (**H**) Quantification of the colocalization observed in (**G**). Mander’s coefficient M2 quantified only sites with *Trp53inp2* signal. Ordinary one-way ANOVA p < 0.0001. Tukey’s multiple comparisons tests: Control vs -NGF p=0.0227; Control vs +NGF p=0.0017; -NGF vs +NGF p<0.0001. n=5

All RNA functions, including splicing, nuclear export and active intracellular transport, depend on the interaction with RNA-binding proteins (RBPs)^22^. We recently performed a proteomic analysis to identify the RBPs interacting with the 3*’*UTR of *Trp53Inp2* and found that the Transactive response DNA binding Protein 43 (TDP-43) interacted with *Trp53Inp2* 3*’*UTR (M. Darsinou, M.Gaspari and A. Riccio, unpublished results). TDP-43 is a ubiquitous RBP that belongs to the heterogeneous nuclear ribonucleoprotein (hnRNP) family^50^. Although mostly confined to the nucleus where regulates RNA splicing and stability, TDP-43 has also been detected in the cytoplasm in physiological conditions and in neurodegeneration^51,52^. First, we confirmed that TDP-43 was localised in axons in control conditions, and that the expression increased in response to NGF (**Fig. S6E**). In control conditions, TDP-43 was significantly more abundant in the nucleus compared to the cytoplasm. However, no significant differences were observed after NGF stimulation possibly due to the *“*leaking*”* of TDP-43 into the cytoplasm in response to cellular stress associated with NGF deprivation (**Fig.S6F, G**). These experiments indicate that in sympathetic neurons, TDP-43 is localised extranuclearly and that axonal localisation changes in response to NGF.

A recent study showed that TDP-43 may represent a previously unknown m^6^A reader acting as an m^6^A-binding protein^29^. To find whether TDP-43 interacts with *Trp53inp2* in a methylation-dependent manner, we first performed RIP in control and METTL3-inhibited sympathetic neurons. Inhibition of methylation significantly decreased TDP-43 binding to *Trp53inp2* (**Fig.5E**). Similar results were observed when RIPs were performed in distal axons (**Fig.5F**). Given that *Trp53inp2* transport to distal axons increases in response to NGF, we hypothesized that binding to TDP-43 may regulate transcript localisation. Using immuno-FISH, we found that co-localization of TDP-43 and *Trp53inp2* was moderate under control conditions, became stronger during NGF withdrawal, and plummeted after exposure to NGF (**Fig.5G-H**). The timing of the interaction suggests that TDP-43 contributes to *Trp53inp2* transport to distal axons by initially forming a complex in the cell bodies. Once the RNA-RBP complex has reached the distal axons, *Trp53inp2* may be released to bind the RNA-binding protein HuD and the NGF receptor TrkA to mediate NGF signalling^37^.

### m^6^A methylation of Trp53Inp2 promotes neuronal survival and axon growth

In sympathetic neurons, *Trp53Inp2* interacts with the NGF receptor TrkA to increase NGF/TrkA complex internalisation and to promote the retrograde transport of signalling endosomes to the cell bodies^37^. To study whether m^6^A methylation regulates *Trp53Inp2* functions, sympathetic neurons dissected from *Trp53Inp2* floxed mice (*Trp53Inp2*^*fl/fl*^) were grown in compartmentalised chambers for 7 days and infected with adenoviral vectors (AAV) expressing the Cre recombinase. In this model system, sympathetic neurons are seeded in the central compartment and NGF (100 ng/ml) is added to the lateral compartments to allow rapid and extensive axon growth (**Fig.6A**, top left). In neurons lacking *Trp53Inp2* (*Trp53Inp2*^*fl/fl*^*+Cre*) axon growth was remarkably stunted when compared to controls (**Fig.6A**). Growth defects were completely rescued by co-infecting *Trp53Inp2* wild type (*Trp53Inp2*^*fl/fl*^*+Cre+Trp53Inp2 WT*, **Fig.6A**, bottom panels). Conversely, in neurons lacking *Trp53Inp2* and infected with an AAV expressing *Trp53Inp2* bearing A695G mutation that cannot ne methylated on an adenosine within the 3*’*UTR, axon growth defects were not rescued (*Trp53Inp2*^*fl/fl*^*+Cre+Trp53Inp2* ^*A695G*^, **Fig.6A**, bottom panels).

**Fig. 6.**
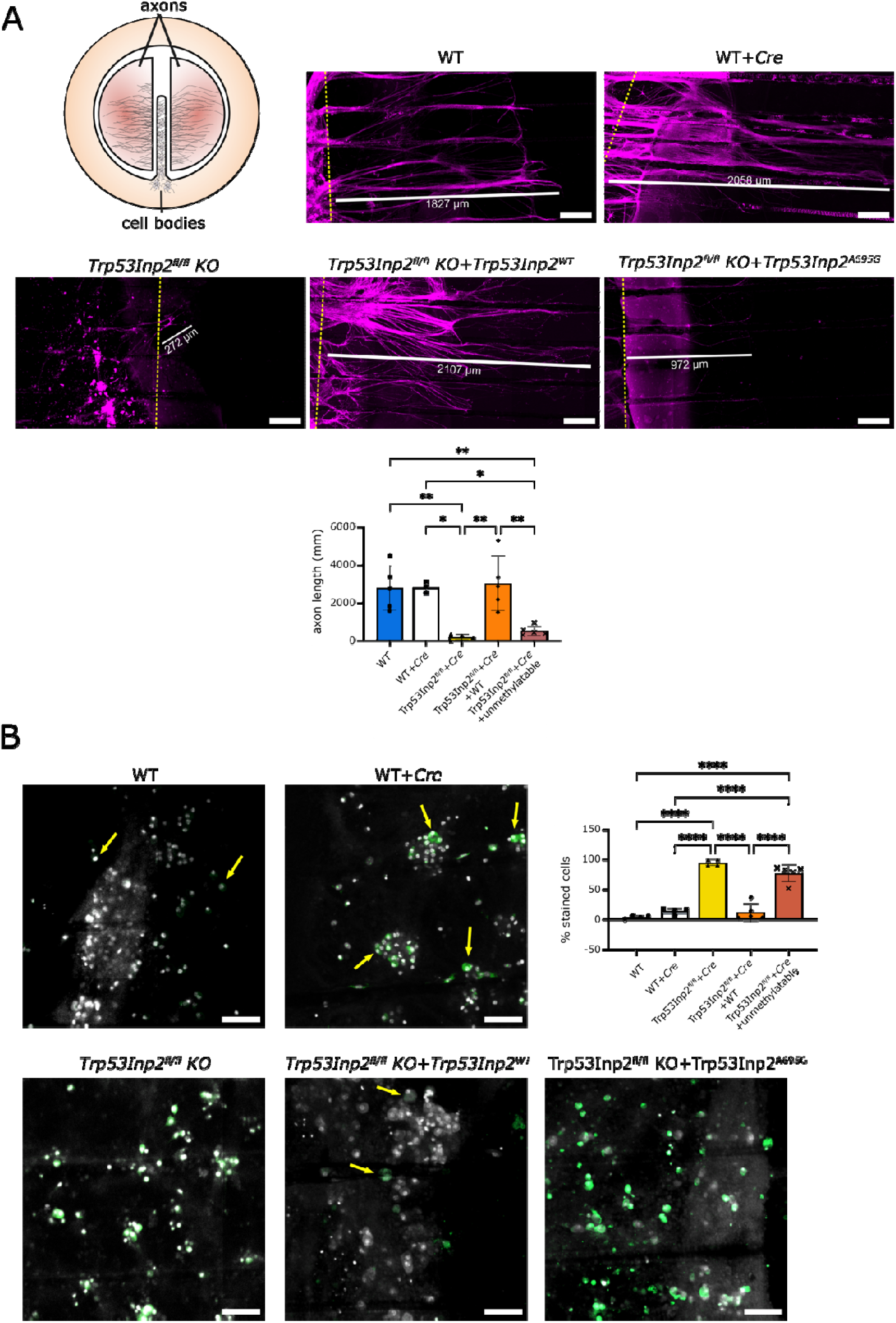
Trp53inp2 3’ UTR methylation is essential for axon growth and neuronal survival. (**A** *left*) In compartmentalised chambers cell bodies are cultured with low NGF (10 ng/ml,) in the central compartment and a higher concentration of NGF (100 ng/ml) in the lateral compartments to promote axon growth. Confocal microscopy images of F-Actin (magenta) to label distal axons under the indicated conditions. Yellow dotted lines indicate the inner wall edge of the chamber. Wildtype neurons (WT); WT neurons infected with Cre AAV (WT+Cre); *Trp53inp2*^*fl/fl*^ neurons infected with Cre AAV (*Trp53inp2*^*fl/fl*^ KO); *Trp53inp2* KO neurons infected with AAV encoding *Trp53inp2* wild-type (*Trp53inp2*^*fl/fl*^ KO+*Trp53inp2*^*WT*^). *Trp53inp2*^*fl/fl*^ KO infected with AAV expressing *Trp53inp2* with an A G mutation (*Trp53inp2*^*fl/fl*^ KO+*Trp53inp2*^*A695G*^). Cultures were infected at DIV7 and grown until DIV14. (*Bottom*) Quantification of axonal length. Ordinary one-way ANOVA, WT vs. *Trp53inp2*^*fl/fl*^ KO, *p*=0.0048; WT vs. *Trp53inp2*^*fl/fl*^+*Trp53inp2*^*A695G*^, *p*=0.0093; WT+Cre vs. *Trp53inp2*^*fl/fl*^ KO, *p*=0.0127; WT+Cre vs. *Trp53inp2*^*fl/fl*^ + *Trp53inp2*^*A695G*^, *p*=0.0247; *Trp53inp2*^*fl/fl*^ KO vs. *Trp53inp2*^*fl/fl*^ KO+*Trp53inp2*^*WT*^, *p*=0.002; *Trp53inp2*^*fl/fl*^ KO+*Trp53inp2*^*WT*^ vs. *Trp53inp2*^*fl/fl*^+*Trp53inp2*^*A695G*^, *p* = 0.0040. (*B*) Confocal microscopy of cell bodies. IT DEAD (green) labels neurons whose membranes have become permeable due to cell death. The outlines of DAPI-stained nuclei are shown in white. *Top-right:* Quantification of degenerated cell bodies. Ordinary one-way ANOVA, WT vs. *Trp53inp2*^*fl/fl*^ KO, *p*<0.0001; WT vs. *Trp53inp2*^*fl/fl*^+*Trp53inp2*^*A695G*^, *p*<0.0001; WT+Cre vs. *Trp53inp2*^*fl/fl*^ KO, *p*<0.0001; WT+Cre vs. *Trp53inp2*^*fl/fl*^+*Trp53inp2*^*A695G*^, *p*<0.0001; *Trp53inp2*^*fl/fl*^ KO vs. *Trp53inp2*^*fl/fl*^ KO+*Trp53inp2*^*WT*^, *p*<0.0001; *Trp53inp2*^*fl/fl*^ KO+*Trp53inp2*^*WT*^ vs. *Trp53inp2*^*fl/fl*^+*Trp53inp2*^*A695G*^, *p*<0.0001.

The interaction of *Trp53Inp2* with TrkA receptors promotes retrograde transport of the signalling endosomes to the cell bodies, an event necessary for neuronal survival. To study whether *Trp53Inp2* m^6^A methylation was necessary for sympathetic neuron survival, neurons were co-infected either with *Trp53Inp2*^*fl/fl*^*+Cre* and *Trp53Inp2*^*WT*^ or *Trp53Inp2*^*A695G*^. In neurons lacking *Trp53Inp2* we observed remarkable cell death that was not rescued by expressing *Trp53Inp2*^*A695G*^ (**Fig. 6B**). Conversely, infection of *Trp53Inp2*^*WT*^ substantially rescued neuronal survival. Together these data indicate that m^6^A methylation of *Trp53Inp2* is essential for the non-coding functions of the transcript and for mediating axon growth and survival of NGF-dependent developing sympathetic neurons.

## Discussion

Epigenetic modifications of RNA are emerging as an essential mechanism that regulates RNA metabolism. Here we show that upon rapid stimulation of sympathetic neurons with the neurotrophin NGF, many m^6^A methylated transcripts are rapidly transported to distal axons and growth cones where they are stored mostly untranslated. We also discovered that methylation of the bi-functional mRNA *Trp53Inp2* is necessary for maintaining translation inhibited and for supporting axon extension and cell survival.

### NGF induces the rapid transport of methylated transcripts in axons

Localisation of RNA in axons and dendrites is necessary for axon extension during development^5,11,13^ and for supporting nerve regeneration^6,53^, dendritic plasticity and synaptic functions in adulthood^7,54-56^. Many factors contribute to the sorting and peripheral transport of transcripts, including localisation elements mostly present within the 3*’*UTR^3,13,17^. Axonal RNAs express longer and more complex 3*’*UTRs harbouring multiple localisation motifs that recruit RNA binding proteins (RBPs), whose combinations contribute to determining transcript localisation^3,10,12^. Here we demonstrate that epigenetic modification of the RNA m^6^A plays a key role in mediating axonal transport. m^6^A is highly abundant in nerve cells and especially enriched at the 3*’*UTRs where it regulates many aspects of RNA metabolism including RNA splicing, nuclear export and translation^23,25^. However, we provide the first evidence that m^6^A modifications plays a key role in inducing the rapid localisation of transcripts in axons in response to an extracellular signal. We discovered almost 800 transcripts hypermethylated in axons shortly after NGF stimulation (46% of axon-enriched RNAs in all conditions) and found that these RNAs encode proteins involved in cell-to-cell signalling (**Fig. 2D**). A previous study has shown that in sensory neurons NGF induces demethylation of axonal RNAs by activating the demethylase FTO in axons, and this event is essential for the translation of an axonal transcript^57^. Because in sympathetic neurons the m^6^A writers (METTL3, METTL14 and WTAP) and erasers (FTO and ALKBH5) are not expressed outside the nucleus, NGF mostly induces the rapid mobilization of methylated transcripts from the cell bodies to axons. Interestingly, in homeostatic conditions when NGF is supplied for longer times, the levels of m^6^A methylated transcripts in axons return to levels similar the ones observed in control conditions. Together these data suggest that degradation of methylated RNA takes place within hours and is consistent with recent data showing that YTHDF proteins acts redundantly to mediate RNA decay^58^. Given that NGF signalling activates the ERKs and PI3K pathways, it is possible that NGF-induced posttranslational modifications of RBPs may regulate the interaction with m^6^A methylated elements or their assembly into complexes that increase RNA transport in axons.

### m^6^A methylation inhibits the translation of axonal Trp53Inp2

Ribo-Tag analysis of RNAs engaged with ribosomes revealed that in response to a short exposure to NGF only a handful of transcripts were translated. Conversely, both RNA methylation and translation remained largely unchanged in cell bodies. m^6^A modification has been shown to both promote protein synthesis in the adult regenerating sciatic nerve^59^, and to decrease activity-dependent translation in cortical neurons^60^. In sympathetic neurons, methylation mostly maintain transcripts untranslated during the transport to distal axons and growth cones. This is an essential mechanism ensuring that transcripts do not undergo premature translation and potential mRNA decay while in transport. In addition, m^6^A methylation acts as a *“*translational switch*”* for bifunctional mRNAs harbouring both coding and non-coding functions. There are increasing evidence that many mRNAs may function as structural or scaffolding non-coding RNAs in certain cell types or under certain conditions, while being translated in others^46,61^. Our meRIP-Seq revealed that *Trp53Inp2* is amongst the most actively transported m^6^A-RNAs in axons in response to NGF. *Trp53Inp2* mRNA is translated in an autophagy protein in myocites^47^, but in sympathetic neurons, interacts with the NGF receptor TrkA to promote NGF signalling, axon growth and neuronal survival^37^. Inhibition of m^6^A methylation inhibits *Trp53Inp2* axonal transport in neurons and promotes translation, acting as a switch between coding and non-coding functions. This finding is especially relevant in developing neurons where the epitranscriptome undergo substantial changes in response to trophic signals and guidance molecules. Thus, changes of RNA methylation provide an additional mechanism regulating not only protein synthesis but also mRNA non-coding functions.

### How are m^6^A methylated mRNAs translated in axons?

Our data support a model by which m^6^A methylated mRNA is transported to distal axons and growth cones in response to acute stimulation with NGF. However, NGF promotes protein synthesis in axons of developing neurons, and we discovered that prolonged exposure to NGF results in decreased transport of axonal methylated transcripts. There are several potential explanations of these results. First, it is possible that when axons are deprived of NGF and restimulated, the first response is to methylate critical RNAs to ensure that transcripts are rapidly transported to distal axons and growth cones to be locally translated (**Fig.1A-C**). Following prolonged NGF exposure, neurons reach a homeostatic state whereby the continuous transport of mRNAs is balanced by translation and decay, as well as other forms of transport. A second contributing factor may be that the first wave of methylated mRNAs may not be translated and act as non-coding, structural RNAs. The finding that *Trp53Inp2* and other long non-coding RNA are enriched in cortical neuron dendrites, are transported in axons in response to NGF suggests that the first wave of methylated mRNAs reaching the distal axons remain preferentially untranslated, possibly holding coding-independent functions aimed at enhancing NGF signalling. Finally, the meRIP-Seq revealed that m^6^A is enriched in the 3*’*UTRs of axonal transcripts. It has been widely reported that neuronal RNAs tend to have an increased average number of m^6^A bases^60,62^, but it is unclear whether they act redundantly or if each base affects RNA differently. For example, we discovered that mutation of one of the three m^6^A within *Trp53Inp2* 3*’*UTR is sufficient to affect axonal growth. We also previously discovered that the in sympathetic neurons, 3*’*UTR may undergo cleavage in a process that increases the translational efficiency of the mRNAs expressing a shorter 3*’*UTR, and at the same time generating a new class of non-coding RNAs with yet unknown functions^10,12^. Further studies confirmed that 3*’*UTR cleavage is a remodelling process that takes place in many cell types and across the species ^63-66^. It is conceivable that at least for some mRNAs, the cleavage of the 3*’*UTR removes the methylated bases that cause translational inhibition, allowing the translation of transcripts expressing a shorter 3*’*UTR. Further experiments will be needed to study the correlation between 3*’*UTR cleavage and m^6^A methylation.

### Does m^6^A methylation provide a link between neurodevelopment and neurodegeneration?

Similarly to DNA, epigenetic modifications of the RNA rely on writers, erasers and readers. While writers and erasers add or remove the epigenetic marks respectively, readers bind to the methylated elements and play a key role in translating m^6^A modifications into specific RNA functions ^24^. Several m^6^A readers have been identified so far, including the YTH family protein^67^, the Heterogeneous Nuclear Ribonuclear Protein C (HNRNPC)^28^ and in neurons, the Fragile X Mental Retardation Protein (FMRP ^32,33^ and the Transactive response DNA binding Protein (TDP-43) ^29^, a predominantly nuclear protein that binds to RNA transcripts and mediates splicing, nuclear export and stability. Mis-localization and mutation of TDP-43 have been linked to Amyotrophic Lateral Sclerosis (ALS) a severe neurodegenerative disorder of the motoneurons^68^. We found that upon exposure to NGF, TDP-43 localises to the cytoplasm. Recent studies have indicated that TDP-43 bind to methylated transcripts and that RNAs are hypermethylated in the spinal cord of ALS patients ^29^. We discovered that TDP-43 is among the RBPs that bind in axons to the 3*’*UTR of *Trp53Inp2* in a methylation-dependent manner and that the binding increases in response to NGF withdrawal, when the complexes are detected in distal axons. An intriguing interpretation of these results is that m^6^A methylation of axonal RNAs plays a key role in axon growth and potentially in response to nerve injury, providing a new exciting link between mechanisms of neurodevelopment and neurodegeneration.

### Experimental procedures

#### Cell lines

C2C12 cell lines were purchased from ATCC and were cultured in DMEM supplemented with 10% fetal bovine serum, 2 mM L-glutamine, and 1X Penicillin-Streptomycin. For differentiation, cells were grown to 60% confluency, washed with PBS, and cultured with DMEM supplemented with 2% horse serum, 2 mM L-glutamine, and 1X Penicillin-Streptomycin. Medium was changed every 2 days until day 7. AD293T cells for virus production were purchased from ATCC and cultured in DMEM supplemented with 5% fetal bovine serum, 2 mM L-glutamine, and 1X Penicillin-Streptomycin.

#### Sympathetic neuron cultures

All animal studies were approved by the Institutional Animal Care and Use Committees at University College London. All cell culture reagents were sourced from Thermo-Fisher. Superior cervical ganglia (SCG) were dissected from post-natal day 0-2 C57BL6/J wildtype mouse pups of both sexes and then used for explant cultures or enzymatically dissociated and grown on coverslips or compartmentalised cultures, as previously described ^69^. Cultures were maintained in DMEM media supplemented with 10% foetal bovine serum, 2 mM L-glutamine, 1X Penicillin-Streptomycin, 10 µM cytosine arabinoside (AraC), and 50 ng/ml Nerve Growth Factor (NGF). Explants and dissociated cultures were grown on 12 mm coverslips coated with rat-tail collagen and 5 ng/ml laminin for 4-8 days, as indicated.

When performing experiments involving NGF starvation, cells were washed twice with PBS and 50 µM BAF caspase inhibitor (Cayman Chemical, Cat. 16118) was added to medium without NGF. Neurons were cultured for 18 hours and then either used for experiments or restimulated by adding NGF to the medium.

For metondylation inhibition experiments, cells were cultured as normal and METTL3 inhibitor STM2457 (MedChemExpress, Cat. HY-134836) was added to the medium at a final concentration of 10 µM at the time of plating or at DIV4.

#### mRNA isolation and m^6^A dot-blot

To isolate mRNA, Dynabeads mRNA DIRECT Purification Kit (TermoFisher Scientific, Cat. 61011) were used following the manufacturer*’*s instructions. In brief, cells were washed with 1xPBS and pelleted at 500 g for 5 minutes at 4^°^C. 300 µl of Lysis Buffer were added to the pellet, which was resuspended with 30 seconds of vigorous pipetting. 50 µl of Dynabeads Oligo(dT)_25_ was washed thrice with Lysis Buffer. The cell lysates were added to the washed beads and mixed gently. The mixture was incubated for 5 mins at room temperature on a rotator to allow the binding of polyA tails to the beads. The tubes were then placed on a magnetic rack until clear, and the supernatant was discarded. Beads were washed in 600 µl of Washing Buffer A and 300 µl of Wasing Buffer B. mRNA was eluted by resuspending the beads in 10 µl of Elution Buffer followed by 2 minutes incubation at 80°C. Tubes were immediately placed in a magnetic rack and the supernatant containing mRNA was moved to a new tube.

For dot-blot experiments, mRNA concentration was measured and tubes with 50 ng, 25 ng, and 5 ng in 5 µl volumes were prepared for each sample. mRNA was denatured at 95°C for 3 mins and chilled immediately. A Hybond-N+ membrane (Cytiva, Cat. RPN303B) was divided in equal parts and labelled with pencil. mRNA droplets of 5 µl were added directly onto the membrane and let dry before being crosslinked twice with a Stratalinker 2400 UV Crosslinker in auto mode. Membranes were washed twice in 0.02% Tween-20 in 1XPBS before being blocked for 1 hour at room temperature in 1% BSA in washing buffer. Membranes were incubated overnight at 4°C with anti-m^6^A antibody (ab151230, 1:1,000), washed 3 times for 5 mins, and incubated for 1 hour with goat anti-rabbit HRP-conjugated antibody (1:20,000). Membranes were washed 4 times for 10 minutes with gentle shaking and developed with ECL using the Alliance Q9 Advanced system (UviTec). Images were quantified using ImageJ.

#### Western blotting

Unless otherwise stated, total protein was isolated using RIPA lysis buffer and quantified using the Pierce BCA Protein Assay (ThermoFisher, Cat. 23225). 30 µg of protein per sample was resuspended in LDS Sample Buffer (ThermoFisher, NP0007) and 2.5% *β*-mercaptoethanol and separated by electrophoresis in a NuPAGE Bis-Tris 4-12% mini-gel on a Mini Gel Tank (ThermoFisher, A25977). After transfer to a PVDF membrane, membranes were blocked with 5% milk in TBST. Primary immunoblotting of the membranes was performed overnight at 4^0^C followed by washing and incubation with species-appropriate secondary antibody. Membranes were developed using SuperSignal ECL (ThermoFisher, Cat. A38554).

#### Immunocytochemistry

Cells were washed with PBS, fixed using 4% paraformaldehyde for 10 minutes, followed by permeabilization using 0.2% Triton X-100 for 10 minutes and three washes. Cells were blocked in 1% BSA for 1 hour at room temperature and then incubated with primary antibodies (Table S1) in blocking buffer for 1.5 hour at room temperature or overnight at 4°C (depending on the antibody). Cells were washed with PBS 3 times for 5 minutes and incubated with the species-appropriate secondary antibody (1:1,000) for 45 minutes at room temperature protected from light. If the experiment required DAPI (1:1000) or Phalloidin (1:1000) staining, the reagents were added to the secondary antibody solution. Cells were washed 3 times for 5 minutes in PBS and finally mounted on glass slides using Fluoromount-G mounting medium (Fisher Scientific, Cat. 15586276). Slides were incubated for at least 4 hours and imaged on an LSM 900 Airyscan 2 confocal microscope (Zeiss Microscopy) using 10x Air EC Plan Neofluar (NA: 0.3), 40x Oil Plan Apochromat (NA: 1.3), and 63x Oil Plan Apochromat (NA: 1.4) objectives and Immersol 518F immersion oil.

#### RNA immunoprecipitation

RNA immunoprecipitation was performed as previously described ^37^. Briefly, cells were washed with PBS and lysed using lysis buffer (150 mM NaCl, 50 mM Tris-HCl, ph 8.0 1.5% Triton X-100, proteinase inhibitor cocktail, 200 U/ml RiboLock RNAse Inhibitor). Lysates were centrifuged at 200 g for 10 minutes at 4°C and supernatants were used in the next steps. 10% was saved as input. Dynabeads Protein G (Invitrogen, Cat. 10003D) were resuspended and washed 3 times in lysis buffer. The mixture was rotated for 1 hour at 4°C for clearing. After separating the magnetic beads, the supernatants were transferred to pre-chilled tubes and 5 µg of TDP-43 or IgG isotype control antibody were added followed by overnight rotation at 4°C. The next day, 40 µl of beads were washed and combined with the lysates, rotated for 2 hours at 4°C followed by extensive washing 4 times for 10 minutes at 4°C. On the final wash, the supernatants were transferred to fresh tubes to reduce non-specific binding and beads were separated on a magnetic rack. Supernatants were discarded and beads resuspended in 100 µl lysis buffer. 350 µl of LB from the PureLink RNA Mini Kit (Invitrogen, Cat. 12183018A) containing 2.5% *β*-mercaptoethanol was added to inputs and immunoprecipitated samples. After vigorous vertexing, beads were incubated 5 minutes for RNA elution and separated using a magnetic rack. Supernatants were collected, and RNA was purified following the PureLink RNA Mini Kit*’*s instructions for RNA clean-up. All samples were DNAse digested using the TURBO DNAse kit according to the manufacturer*’*s instructions (Invitrogen, Cat. AM2238).

#### RT-qPCR

RNA was isolated from neurons using PureLink RNA Mini Kit and reverse transcribed using the RevertAid cDNA synthesis kit (Invitrogen, Cat. K1622). RT-qPCR was performed with Luna Universal qPCR Master Mix (New England Biolabs, Cat. M3003S) following the manufacturer*’*s standard protocol, and 2 µl of cDNA. Reactions were set up in triplicates and run on QuantStudio 6 Pro machines. Data were analysed using the comparative Ct method (ΔΔCt). Melting curves and no template controls were included to verify primer and reaction specificity, respectively. Primer sequences are described in supplementary table S2.

#### Compartmentalised cultures

For viral transduction experiments, 35 mm dishes were coated with rat-tail collagen and 5 ng/ml laminin, air-dried, and scratches were made on the collagen. Compartmentalised chambers were seated onto the dish using silicon vacuum grease, left to dry overnight, and 100,000 neurons were plated in the central compartment^13^. Cultures were maintained in DMEM media supplemented with 10% foetal bovine serum, 2 mM L-glutamine, and 1X Penicillin-Streptomycin, and 50 ng/ml Nerve Growth Factor (NGF) until DIV7, after which the central compartment was reduced 10 ng/ml NGF and the side compartments maintained at 50 ng/ml until DIV14. For all other experiments requiring axonal fractions (including m^6^A-sequencing), 2 million dissociated neurons were cultured on each transwell membrane insert (Corning, Cat. 3414) and cultured in the medium described above until DIV 4, when NGF was only provided to the bottom compartment to promote directional axonal growth. At DIV15, subcellular fractions were collected as described by Flamand & Meyer^70^.

### m^6^A RNA Immunoprecipitation (meRIP) and axon-meRIP-seq

meRIP conditions were adapted from^60^ following a previously published protocol for low-input samples^71^. Briefly, Dynabeads Protein G magnetic beads were washed and resuspended in 250 µl reaction buffer (150 mM NaCl, 10 mM Tris-HCl, pH 7.5, 0.1% NP-40, 200 U/ml RiboLock RNAse inhibitor). 5 µg of m^6^A or IgG control antibodies were added, and the mixtures were rotated overnight at 4°C. The next day, magnetic beads were separated and washed 3 times with washing buffer (50 mM NaCl, 10 mM Tris-HCl, pH 7.5, 0.1% NP-40) before resuspending them in 250 µl reaction buffer. Samples were spiked with 0.1 fmol of *in vitro* transcribed FluC unmodified RNA as a negative control. 10% of purified RNA was saved as input and the remaining RNA (up to 250 µg) was added to the resuspended beads. Mixtures were rotated for 2 hours at 4°C. Beads were separated magnetically, the supernatants were discarded, and the beads were washed 3 times for 10 minutes. For the final wash, samples were moved to RNAse-free tubes and beads were separated and resuspended in 350 µl LB buffer from the PureLink RNA Mini Kit. To elute the RNA, samples were vigorously vortexed and incubated for 5 minutes. Beads were separated and the RNA-containing supernatants were purified following the kit*’*s RNA clean-up protocol.

For axon meRIP-Seq, neurons were NGF-treated as indicated and RNA was isolated from either axons or cell bodies using the PureLink RNA Mini Kit. After saving 10% RNA for inputs, RNA was sheared into ∼200 bp fragments using the NEBNext® Magnesium RNA Fragmentation Module (New England Biolabs, Cat. E6150S). After purification with AMPure XP beads (Beckman Coulter, Cat. A63880), meRIP was performed as described above. RNA size, quality, and concentration were verified via TapeStation High Sensitivity RNA ScreenTape Analysis. Using the Watchmaker RNA Library Prep Kit with Polaris Depletion (Watchmaker Genomics, Cat. 7BK0002-024), rRNA and Goblin RNA depletion, first and second strand cDNA synthesis, sequencing library prep and amplification were performed in one continuous workflow. Libraries were QC*’*d and sequenced according to the manufacturer*’*s instructions using Illumina NextSeq 2000 platform (100 cycles) to generate ∼20-40 million 50 bp paired-end reads.

### Translating Ribosome Affinity Purification and sequencing (TRAP-Seq)

Sympathetic neurons from B6J.129(Cg)-Rpl22tm1.1Psam/SjJ (Jackson Laboratory, Strain #029977) RiboTag x DBH-Cre ^72^ were dissected at P0-P2 and cultured on transwell inserts under the indicated conditions. The TRAP protocol was performed as described ^36^. Briefly, cells were treated for 15 minutes with cycloheximide, then both sides of the transwell inserts were washed 3 times with ice-cold PBS + 100 µg/ml cycloheximide solution. Axons and cell bodies were collected as described above, and pellets were resuspended and mechanically lysed by vigorous pipetting in 1 ml of fresh CHX-Lysis Buffer (20 mM HEPES-KOH, 5 mM MgCl_2_, 150 mM KCl, 1 mM dithiothreitol (DTT), 1% NP-40, 100 µg/ml cycloheximide, 200 U/ml RiboLock RNAse inhibitor, EDTA-free proteinase inhibitors cocktail). 10% of lysates were saved as input and the remaining were transferred to tubes with washed Dynabeads Protein G and cleared by rotating them for 1 hour at 4°C. Beads were separated and the lysates were transferred to a new tube with 2.5 µg of HA antibody and rotated overnight at 4°C. The next day, fresh beads were washed, mixed with the lysate and antibody solution, and rotated for 2 hours at 4°C. Beads were separated and washed 3 times for 10 minutes in washing buffer (20 mM HEPES-KOH, 5 mM MgCl_2_, 350 mM KCl, 1 mM dithiothreitol (DTT), 1% NP-40, 100 µg/ml cycloheximide). For the final wash, samples were transferred to RNAse-free tubes, and the beads were separated and resuspended in 350 µl LB buffer from the PureLink RNA Mini Kit. To elute the RNA, samples were vigorously vortexed and incubated for 5 minutes. Beads were separated and the RNA-containing supernatant was purified following the kit*’*s RNA clean-up protocol. RNA size, quality, and concentration were verified via TapeStation High Sensitivity RNA ScreenTape Analysis. Using the NEBNext® Single Cell/Low Input RNA Library Prep Kit for Illumina® (New England Biolabs, Cat. E6420S), first and second strand cDNA synthesis, sequencing library prep and amplification were all performed in one continuous workflow. Libraries were QC*’*d and then sequenced according to the manufacturer*’*s instructions using Illumina NextSeq 2000 platform (100 cycles) to generate ∼30 million 101 bp single-end reads.

### Sequencing data analysis

Sequencing data was processed using nf-core/rnaseq (https://zenodo.org/records/7998767) of the nf-core collection of workflows ^73,74^. Reads were pre-processed including quality filtered and trimmed before being mapped to GRCm38, using a combination of fastqc, samtools, star, salmon, trimgalore (full list of software and processes in table S3) set with default parameters. Mapped reads were de-duplicated and quantified to produce gene counts which were used as input for differential gene expression. Differential gene expression was performed with the SARTools pipeline ^75^ using DESeq2 ^76^. Differential gene expression analysis was performed using edgeR ^77^ using TMM normalization. Only genes with more than 15 total counts with at least 10 counts in one of the samples were considered for further processing. Surrogate Variable Analysis was used to remove batch effects ^78^. Linear models were fitted through all eight conditions (∼ SV1 + SV2 + compartment * seqmethod * treatment). Gene Ontology analysis was performed using Metascape ^79^ and plots were created with ggplot2 (Wickham, 2009) on R (R Core Team, 2014).

### HCR RNA-FISH and immuno-FISH

HCR probe sets targeting the *Trp53inp2* (B1 initiator, 30 split-initiator probes) and *Lin7c* (B3 initiator, 30 split-initiator probes) were purchased from Molecular Instruments. Experiments were performed based on the manufacturer*’*s protocol for mammalian cells ^80^. All reagents and materials used were RNAse-free. Transfected cells were fixed with 4% paraformaldehyde (TAAB) at room temperature for 10 minutes followed by permeabilization in 70% ethanol for 3 hours at 4°C. Cells were washed 2 times for 5 minutes in 2x SSCT Buffer (2x SSC +0.1% Tween20) and pre-hybridized in Probe Hybridization Buffer for 30 minutes at 37°C. Cells were incubated with 1.2 pmol of each probe overnight at 37°C in a humidified chamber. Excess probes were washed with Probe Wash Buffer 4 times for 5 minutes at 37°C, followed by 2 washes for 5 minutes in 5x SSC Buffer at room temperature. Pre-amplification was performed in Amplification buffer for 30 minutes at room temperature. 18 pmol of each fluorescent hairpin amplifier (B1h1/B1h2 Alexa Fluor 647 and B3h1/B3h2 Alexa Fluor 594) were snap cooled in separate tubes by heating them for 90 seconds at 95°C in a pre-warmed thermocycler and allowed to cool in the dark for 30 minutes. After pre-amplification, the buffer was removed from the cells and replaced with cooled hairpins mixed in Amplification Buffer. For quantitative HCR imaging, amplification was performed for 45 minutes in the dark at room temperature. Excess hairpins were washed in 5x SSCT Buffer 5 times for 5 minutes, followed by 10 minutes incubation in 1 μg/mL DAPI in 1x PBS. Cells were mounted in Pro-Long Gold antifade mountant (#P36930, Thermo Fisher Scientific) and incubated overnight. Negative controls without probes and without amplification were included for each experiment. For immuno-FISH, cells were fixed with 4% PFA for 10 minutes in the dark and a standard immunocytochemistry protocol was followed.

Airyscan imaging was performed using a Zeiss LSM900 confocal microscope with a 63× Plan Apochromat objective (NA = 1.4) and Airyscan 2 detector with GaAsp technology. Airyscan optimal settings were used for capture, and images were processed using the Zen Blue 3.2 Airyscan 3D processing module with standard settings.

### Puro-Proximity Ligation Assays (Puro-PLA)

Detection of nascent translated proteins was carried out using a proximity ligation assay, where a puromycin antibody was combined with protein-specific antibody. Duolink detection reagents (Sigma-Aldrich, Cat. DUO92008) were used to detect the reaction according to the manufacturer*’*s recommendations and established protocol ^81^. 10 μg/ml puromycin (Sigma-Aldrich, P7255) was added to sympathetic neurons for 15 minutes at 37°C in a humidified chamber with 10% CO_2_. For translation inhibition controls, neurons were treated with various concentrations of anisomycin (Sigma-Aldrich, A9789) or cycloheximide (Sigma-Aldrich, C1988) for either 15 or 30 minutes before incubation with puromycin. Incubation was stopped by three rapid washes in cold PBS, followed by fixation for 10 minutes in 4% Formaldehyde (Thermo Fisher Scientific, 28908) at room temperature. Neurons were permeabilised for 10 mins in 0.02% Triton X-100 in 1X PBS, washed, and blocked for 1 hour in 1% BSA. Primary antibodies were then diluted in 1% BSA and cells were incubated in this dilution for 90 minutes. After three washes, neurons were incubated with anti-rabbit Plus (Sigma-Aldrich, DUO92002) and anti-mouse Minus (Sigma-Aldrich, DUO92008) probes diluted 1:5 in PBS for 1 hour at 37°C. Cells were washed with washing buffer A (0.01 M Tris, 0.15 M NaCl, 0.05 % Tween20) and incubated for 30 minutes with the ligation reaction mix in a humidified chamber at 37°C. After 3 further washes with washing buffer A, cells were incubated with the amplification reaction mix in a humidified chamber at 37°C for 1 hour and 40 minutes. Amplification was terminated by three washes in 1X washing buffer B (0.2 M Tris, 0.1 M NaCl, pH 7.5) followed by a wash in 0.01X wash buffer B. Neurons were stained with fluorescent markers [Ph647 (Thermo Fisher Scientific, A22287), DAPI (Sigma-Aldrich, 268298)] for 1 hour at RT. Finally, the coverslips were washed twice with PBS and twice with Milli-Q water, mounted with Fluoromount G and imaged or stored at 4°C.

### Image analysis

Images captured in Airyscan mode were first Airyscan processed on Zen 3.2 Blue Edition and maximum intensity orthogonal projections were generated. To quantify the distribution of m^6^A RNAs across axons as a function of distance from the axonal tip, background was reduced and the Find Maxima tool in ImageJ was used to record all positions. The distance from the edge was calculated using the Pythagorean theorem and m^6^A RNAs were binned into 20 µm segments. Co-localisation of immuno-FISH images was calculated by masking images using a cytoskeletal channel followed by background reduction and using the Coloc2 plugin in ImageJ. Mander*’*s or Pearson*’*s coefficients were recorded and used for statistical testing. Axon length was quantified using NeuronJ plugin. For toxicity assay, cell death was calculated by masking the cytoskeletal channel and calculating percentage of cells in which there is signal from the toxicity stain.

For HCR RNA FISH, masking using the cytoskeletal channel was performed on maximum projections with a single macro for all images. Afterward, RNA spot quantification was performed using batch processing in the FISH-Quant plugin ^82^ on ImJoy, a hybrid computing platform for deep learning image analysis, with filter sigma = 1.0 and spot detection threshold set at 50.

Dot blots and western blots were quantified by measuring the integrated density grey values in ImageJ. In the case of western blots, this value was then normalised to the loading control.

### In vitro transcription of m^6^A-modified and unmodified RNA

RNA was *in vitro* transcribed from plasmid DNA templates using the HiScribe T7 High Yield RNA Synthesis Kit (New England Biolabs, E2040S) as per the manufacturer*’*s instructions. To produce m^6^A-modified RNA, 10 mM N6-methyl-ATP (Jena Bioscience, NU-1101S) was used and the ratio of ATP: m^6^ATP was changed accordingly, following the manufacturer*’*s protocol for RNA synthesis with modified nucleotides. *In vitro* transcribed RNA was capped using the *Vaccinia* capping enzyme and mRNA Cap 2’-O-methyltransferase (New England Biolabs, Cat. M2080 and M0366) in a one-step reaction according to the manufacturer*’*s protocol. RNAs were polyadenylated using *E. coli* Poly(A) Polymerase (New England Biolabs, Cat. M0276) for 30 minutes following the manufacturer*’*s protocol. RNAs were cleaned up and length and yield were calculated using TapeStation RNA Screentape Analysis.

### siRNA-mediated knockdown and magnetofection

SMARTpool ON-TARGETplus siRNA targeting *Mettl3, GAPD*, as well as a non-targeting pool were purchased from Horizon Dharmacon. siRNA was transfected into C2C12 cells through Neuromag magnetofection (OZ Biosciences, Cat. NM50200) following the Neuromag protocol. Briefly, 50 nM siRNA and 0.5 µg pBIRD GFP reporter were diluted in serum-free medium. Neuromag solution was diluted in a second tube with serum-free medium and both tubes were mixed and incubated for 15 minutes to form complexes. The mixture was added to cells and placed in a 37°C incubator on a magnetic plate. After 30 minutes, the magnetic plate was removed, and cells were cultured for 72-96 hours.

### Cloning and adenovirus production

Full-length *Trp53inp2* was synthesised in a pUC57 plasmid backbone purchased from Genscript. To generate Adenoviral vectors, the AdEasy Adenoviral Vector system (Agilent, Cat. 240009) was used following the manufacturer*’*s protocol. Briefly, *Trp53inp2* was cloned into pCMV-Shuttle vector and linearised with PmeI restriction enzyme. Linearised plasmid was co-transformed via electroporation into BJ5183 competent cells alongside pAdEasy-1 vector. Colonies were screened for recombinant adenoviral plasmid via kanamycin selection and plasmid sequencing. After identification and amplification of successful clones, recombinant adeno-CMV-*Trp53inp2* plasmid was digested with PacI enzyme and magnetofected into AD293T cells for viral production. After 10 days, primary vial stocks were collected through four rounds of freeze/thaw cycles. Primary stocks were then used for viral amplification in AD293T cells and stocks collected via freeze/thaw cycles. Stocks were aliquoted and stored at -80°C. Viruses were titrated using the AdEasy Viral Titer kit (Agilent, Cat. 972500) following manufacturer*’*s instructions. To generate adenoviruses to express non-methylatable *Trp53inp2*, QuikChange II site-directed mutagenesis kit was used (Agilent, 200523) on pCMV-Shuttle-*Trp53inp2* and then the same protocols were followed.

### Adenovirus-mediated knockout and rescues

For functional studies of non-methylatable *Trp53inp2*, sympathetic neurons dissected from P0-P2 *Trp53inp2*^*fl/fl*^ mice were grown in compartmentalised Campenot chambers. On DIV7, neurons were infected with Adeno-Cre-GFP, Adeno-*Trp53inp2*, and/or Adeno-mutant-*Trp53inp2* as indicated. Cell body and axon compartments were thereafter cultured in 10 or 50 ng/ml of NGF, respectively. On DIV14, cells were treated with Image-IT DEAD Green viability stain (ThermoFisher, Cat. I10291) before being fixed and stained with Phalloidin 647 (Santa Cruz, Cat. sc-363797) and DAPI. Cultures were imaged to assess axonal growth and cell death using an inverted LSM900i Airyscan 2 microscope with a 10x Air EC Plan Neofluar (NA: 0.3) objective and Immersol 518F in confocal mode.

### Statistical Analysis

All data presented are represented as means and standard deviations of at least 3 independent biological experiments. Two-tailed t-tests, one-way ANOVA, or two-way ANOVA were used to compare means across groups as indicated in figure legends. A *p-value* ≤ 0.05 was chosen as statistically significant. Analysis and plots were done on GraphPad Prism 10.5.0 software.

### References without DOI

Wickham, H. (2009) ggplot2: elegant graphics for data analysis. Springer New York.

R Core Team (2014). R: A language and environment for statistical computing. R Foundation for Statistical Computing, Vienna, Austria. URL http://www.R-project.org/

## Acknowledgements

We thank R. Kuruvilla (Johns Hopkins University) and W.G. Tourtellotte (Northwestern University) for kindly providing Tp53inp2^fl/fl^ and DBH-Cre mice, respectively. We thank M. Darsinou (LMCB, UCL) for help evaluating *Tp53inp2*^*fl/fl*^ mouse knockout efficiency and G. Modol-Caballero (King’s College London) for help optimizing HCR RNA FISH protocols. We also thank all present and past members of the Riccio and Lloyd labs for helpful discussions and insights. This work was supported by Wellcome Trust Investigator Awards 103717/Z/14/Z and 217213/Z/19/Z (to AR), a collaborative grant between University College London and the Neuroscience Center Zurich (to AR and GS) and the MRC LMCB Core Grant MC/U12266B (to A.R.)

## Author contributions

AR conceived the work and wrote the manuscripts. BMC conceived the work, performed most experiments and help writing the manuscript. SW, YJ, and SRN helped performing the experiments. SG performed the RNA-seq analysis, supervised by PLG and GS.

